# The influence of parasite load on transcriptional activity and morphology of a cestode and its ant intermediate host

**DOI:** 10.1101/2023.02.02.526817

**Authors:** Tom Sistermans, Juliane Hartke, Marah Stoldt, Romain Libbrecht, Susanne Foitzik

## Abstract

Parasites with complex life cycles are known to induce phenotypic changes in their intermediate hosts to increase transmission to the final host. The magnitude of these changes could increase with the number of parasites, which would be beneficial to co-infecting parasites. Yet, adverse effects of high parasite load (i.e., many parasites in a single host) might stress both hosts and parasites (e.g., through an increased immune response). We investigated the consequences of parasite load on the transcriptional activity and morphology of the cestode *Anomotaenia brevis* and its intermediate host, the ant *Temnothorax nylanderi*. We demonstrated that many differentially expressed host genes shifted with parasite load, and their functions indicate a stronger immune response and fight against oxidative stress in heavily infected hosts. The expression of other host genes responded to infection in an all-or-nothing manner, as did the morphology of the host workers. However, the cestodes became smaller when they competed with other parasites for resources from a single host. Their expression profile further indicated shifts in host immune avoidance, starvation resistance and vesicle-mediated transport. In summary, our study reveals clear consequences of parasite load and highlights specific processes and traits affected by this.

## Introduction

Parasite infection often leads to phenotypic changes in the host that appear to benefit the parasite (Poulin & Thomas, 1999). These shifts in morphology, physiology, behaviour or other life history traits are particularly pronounced and well-studied in parasites with complex life cycles that exploit multiple host species (Heil, 2016). Examples include the induction of fruit mimicry in ants by the nematode *Myrmeconema neotropicum* (Yanoviak et al., 2008) and the reduction of predator aversion in rodents as a result of *Toxoplasma gondii* infection (Hari Dass & Vyas, 2014). In general, these phenotypic changes are thought to facilitate transfer from intermediate to definitive hosts (Poulin & Thomas, 1999; Weinersmith & Faulkes, 2014).

A common interpretation in evolutionary parasitology is to consider phenotypic changes in infected hosts as the “extended phenotype” of the parasite, when they are encoded in the parasite’s genome and consequently are subject to selection in the parasite (Dawkins, 1982; Moore, 2002). This view has been criticized in recent years due to a lack of experimental evidence on the consequences of these phenotypic shifts for the fitness of the parasite (Cézilly et al., 2013; 2010; Poulin & Maure, 2015). The change in host phenotype that follows infection may also be a by-product of infection that is not beneficial to either party, or an adaptive compensatory response of the host to limit the negative consequences of parasite exploitation (Lefèvre et al., 2008; Minchella, 1985; Poulin, et al., 1994).

One way to gain deeper insights into the causes of the altered host phenotype upon infection is to study the consequences of parasite load on the phenotype and fitness of both interacting partners. We might detect divergent phenotypic changes in the host depending on who is responsible for said changes (parasite or host). For example, Maksimowich and Mathis (2000) explored aggression of the salamander *Plethodon ouachitae* in response to high parasite load of the chigger mite *Hannemania dunni*. They found that highly infected hosts were less aggressive than lowly infected hosts even though higher aggression was found to be advantageous for the parasite, aiding in its transmission (Maksimowich & Mathis, 2000). In another example, haemoparasite load affected the throat colour of male Iberian lizards (Megía-Palma et al., 2018), informing females about a male’s ability to deal with parasitic infections.

Another aspect that can be investigated through the lens of parasite load is the consequences for the parasite, for which an increased infection intensity can have diverse outcomes. Benefits for the parasite include synergistic effects, as individual parasites need to invest less to manipulate the host or alter its phenotype to facilitate transmission to the final host (Franceschi, et al., 2008; Yan et al., 2018). This could lead to stronger fitness benefits for the parasite, for example in the acanthocephalan parasite *Pomphorhynchus laevis*, which causes its host to leave its hiding places more often under higher parasite load, aiding in transmission to new hosts (Labaude et al., 2020). Negative consequences are more competition among parasites, as multiple individuals need to share their host’s resources (Bush et al., 2000). Furthermore, a higher parasite load could trigger stronger host immune defences that in turn could harm the parasites (Kaunisto & Suhonen, 2013). Finally, high parasite load might cause premature host death, consequently reducing parasite fitness (De Roode et al., 2008). In some systems, these trade-offs could also affect the phenotype of the parasite, which is particularly well-studied in cestodes, where crowding results in the development of fewer proglottids, i. e. reproductive segments (Heins et al., 2002; Holmes, 1961).

To investigate the effects of parasite load, we used the cestode *Anomotaenia brevis* and its intermediate host, the ant *Temnothorax nylanderi* as a model system. Adult individuals of this tapeworm species reproduce sexually in the gut of birds, most commonly woodpeckers (Plateaux, 1972; Trabalon et al., 2000). Their eggs are released in bird faeces, which *T. nylanderi* ants use to feed their larvae and thus infecting them with *A. brevis*. The parasite larvae penetrate the ant’s gut and develop into cysticercoids in the haemolymph. They remain in this larval stage until the ants are consumed by avian predators (Fig. S1). The phenotype of *T. nylanderi* workers is strongly altered following infection. Their cuticle is less sclerotized and pigmented during the pupal stage, leading to adult ants exhibiting a lighter, softer cuticle (Plateaux, 1972). Moreover, infected workers are less active, suffer from muscular dystrophy (Feldmeyer et al., 2016), receive more social attention (Scharf et al., 2012) and live substantially longer than their uninfected nestmates (Beros et al., 2021). A three-year study (Beros et al., 2021) found no difference in survival between infected workers and uninfected queens; the latter can live up to two decades in this species. Many of the physiological changes can be interpreted as being advantageous to the cestode and go together with transcriptomic shifts that might explain some of the altered traits of infected workers (Beros et al., 2021; Stoldt et al., 2021; Trabalon et al., 2000). For example, genes related to muscle development are downregulated in infected workers (Feldmeyer et al., 2016), possibly explaining not only muscle maldevelopment, but also the reduced ability of infected workers to escape when a nest is opened by a predator (Beros et al., 2015). Upregulation of genes related to oxidative stress resistance (Stoldt et al., 2021) might explain their extended lifespan, which in turn could increase the likelihood that infected workers will be preyed upon by the parasite’s final host, the woodpecker.

The consequences of infection by multiple parasite individuals have not been studied in this system, despite a high variance in parasite load. While some ant workers are infected by a single cestode, others can harbour more than 70 cysticercoids in their haemolymph (data from this study, see below). Here, we exploit this variance to gain deeper insights into the complex relationship between *A. brevis* and *T. nylanderi*. We study the consequences of parasite load on the transcriptional activity and morphology of *A. brevis* and *T. nylanderi*. We predict that a higher parasite load should lead to more stress in both host and parasite. For the host, a high parasite burden may lead to higher potential of the parasite to damage the host (De Roode et al., 2008), possibly resulting in smaller body size and greater expression of stress genes in heavily infected individuals. If the parasites are unable to evade host recognition, a high parasite load might also exhibit an enhanced immune response. However, in view of the possibility of host manipulation, a higher parasite load should also lead to more extreme phenotypic changes in the host, which might be to the advantage of the parasite. For example, many parasites together may be better at subverting the typical innate immune response of insects, which includes encapsulation and melanisation, or they may be better at upregulating host genes important for responding to oxidative stress to extend host lifespan. When looking at the parasite, crowding is a well-studied aspect of cestode biology (Heins et al., 2002), including early life stages (Benesh, 2019). Therefore, we predict cysticercoids to be smaller in heavily infected hosts. In terms of transcriptomic shifts, we expect genes involved in host communication to be differentially expressed, most likely leading to downregulation of this function at high parasite loads, as individual cestodes will need to invest less in manipulation. However, we hypothesise that genes involved in communication and competition with other parasites are upregulated because competition between densely packed cestodes is likely to be greater. In addition, we predict an overexpression of genes that aid in nutrient acquisition or even in combating starvation, as parasites that must share their host with several competitors are likely to have fewer resources available to them.

## Material and methods

### Collection and sampling of ants

Colonies of *T. nylanderi* were collected in November 2019 and February 2020 in the Lenneberg Forest near Mainz, Germany (50.011605” N, 8°10’48.6” E). Colonies of this ant species, whose workers are only about 2-3 mm long, include several dozen workers and inhabit dead branches and acorns on the forest floor. From the colonies we found, we selected 25 colonies, 21 containing infected workers (Table S1). Infection was initially determined by the yellow colouration of infected workers, a sign of infection (Scharf et al., 2012). The ants were transferred to slide nests consisting of a Plexiglas cavity between two slides and placed in plastered nest boxes with three chambers (Stoldt et al., 2021). Colonies were maintained at 20+/-1°C and fed with small crickets and honey twice per week. Water was provided ad libitum. The number of infected workers and queens were counted, and colonies were kept under these conditions for six months.

At the end of July 2020, all infected workers and a sample of several uninfected workers per colony were collected and dissected (N of dissected, uninfected workers per colony see Table S1). Furthermore, we dissected five workers per colony from four uninfected colonies. For all ant workers that were dissected, the presence and number of cysticercoid cestode larvae was noted and their diameter was measured. Furthermore, worker head width was measured from eye to eye, as host body size was shown to decrease in *T. nylanderi* (Scharf et al., 2012) and other systems with increasing parasite load (Dingemanse et al., 2009). All measurements were taken under a Leica stereomicroscope 200x magnification using the LAS v4.5 software.

The fat body of the ants was extracted within five minutes of starting the dissection. We focused our transcriptome analysis on this tissue because it serves as a storage organ and plays an important role in the production of proteins for immune defence, fighting aging, and fertility (Arrese & Soulages, 2010; Negroni, et al., 2019). We obtained the fat body by isolating the first cuticular segment of the abdomen and removing the gut and trachea. What remains were fat cells and the cuticular segment itself, which was promptly transferred to Eppendorf tubes containing 50μl of Trizol, frozen on dry ice and stored at -80°C until RNA extraction. The cestodes were removed from around the midgut, where they are usually located in the haemolymph and separately transferred into Eppendorf tubes containing 50μl of Trizol. They were also frozen on dry ice and stored at -80°C until RNA extraction.

We selected seven infected colonies for transcriptomic analyses with infected workers with different parasite loads (Table S1). These colonies contained at least one queen and on average 93 workers (mean 93.1±43.4). Two of the infected colonies were collected in the field in November 2019, the other five in February 2020 (Table S1). Infected colonies contained between 2 and 24 infected workers and the infection rate varied between 3% and 22%. For each infected colony, we sampled the fat body of one highly infected worker (high parasite load; 18+ cysticercoids), one lowly infected worker (low parasite load; 1-6 cysticercoids) and an uninfected worker. For RNA extraction we homogenized the samples in Trizol using a plastic autoclaved micropistil. We then added 25μl of chloroform and centrifuged for 15 minutes. Afterwards the upper aqueous phase was transferred into Eppendorf tubes containing 25μl of 96% ethanol. RNA was isolated using the Qiagen RNeasy mini extraction kit following protocol specifications. Additionally, we extracted the RNA of one worker per uninfected colony (colony size: 95.8±44.8 workers). One of these colonies was taken from the field in November 2019 and the other three in February 2020. Sequencing was conducted by Novogene (Cambridge, UK). They enriched the mRNA using oligo(dT) beads, then fragmented randomly by adding fragmentation buffer. The cDNA was synthesized by using mRNA template and random hexamers primer, after which a custom second-strand synthesis buffer (Illumina), dNTPs, RNase H and DNA polymerase I were added to initiate the second-strand synthesis. After a series of terminal repair, A ligation and sequencing adaptor ligation, the cDNA libraries were completed through size selection and PCR enrichment. Illumina sequencing was conducted after pooling according to the libraries’ effective concentration and expected data volume (Novogene Co., Ltd). We sequenced a minimum of 9Gb per sample, resulting in around 30 million paired-end 150bp reads.

We isolated the RNA of cysticercoids of five highly infected workers (between 18 and 51 cysticercoids, median = 23) and five lowly infected workers (four ants with one cysticercoid, one ant with two) from five colonies, which were also included in the host gene expression analyses. The RNA was isolated using the same protocol as the fat body samples and samples were sent to Novogene for RNA amplification using the SMARTer kit and RNA sequencing. None of our cestode samples had a lower RNA yield than 9ng, meaning we can assume that the amplification was reliable, as SMARTer amplification only becomes less reliable with an RNA yield below 1ng (Palomares et al., 2019). Library preparation and sequencing were done in the same way as described above. We sequenced a minimum of 9Gb per sample, resulting in around 30 million paired-end 150bp reads.

### Morphometric analysis

To investigate the influence of cestode infection on host body size we conducted two comparisons. First, we analysed whether head width varied between uninfected workers (n = 100) and infected workers (n = 275) from infected colonies (n = 21 colonies) using a linear mixed effects model taking colony identity into account. Second, to investigate the influence of parasite load directly, we focused only on infected workers (n = 267) and used a linear mixed effects model, with colony identity as random effect (n = 21 colonies). For the analysis of parasite load on cestode body size, we again used a linear mixed effects model with colony and ant identity as random effects to investigate the impact of cestode number on cestode diameter, including 2980 cestodes belonging to 267 ants from 21 colonies. All models were checked for normal distribution of their residuals and analysed using an ANOVA.

### Differential gene expression analysis

We removed adapters and contaminating sequences from *A. brevis, Homo sapiens, Escherichia coli* and vectors from the ant reads using FastQScreen v0.14.0 (Wingett & Andrews, 2018). Then, the raw reads were trimmed with fastp (Chen et al., 2018) in paired-end mode for reads of 120 bp or more and an N-base cutoff of 15. Subsequently, the quality of the reads was assessed using FastQC (Andrews, 2010). Filtered reads were mapped against the *T. nylanderi* genome assembly (Jongepier et al., 2022) using HISAT2 v2.1.0 (Kim et al., 2015) with the *--dta* parameter in preparation for genome-guided assembly. The resulting BAM files were sorted and indexed using Samtools v0.1.19 (Li et al., 2009) and used to generate a genome-guided transcriptome assembly using StringTie v1.3.6 (Pertea et al., 2015). To extract transcript sequences, gffread v0.11.4 was applied to the merged GTF files. To count gene abundances, we made use of the script prepDE.py (retrieved from StringTie webpage, Pertea et al. 2015). Subsequently, the predicted amino acid sequences of the transcripts were retrieved using TransDecoder v5.5.0 (Haas et al., 2013) and functionally annotated using InterProScan v5.46-81.0 (Jones et al., 2014). Finally, we obtained the KEGG IDs of the amino acid sequences using the online tool of EGGNOG mapper 2.1.8 (Cantalapiedra et al., 2021).

The gene count matrix was analyzed using DESeq2 (Love et al., 2014), fitting a negative binomial distribution. We used a likelihood ratio test (LRT) to compare a model that included parasite load and colony identity as factors, to a reduced model including only colony identity, and gene expression was considered to differ between at least two of the parasite load categories when the Benjamini-Hochberg adjusted p-value was below 0.05 (ant DEGs). Principal component analysis was performed to visualize whether parasite load caused clustering of samples. We ran a cluster analysis on the ant DEGs using DEGreport to identify groups of genes based similarity in their expression patterns. The minimum cluster size was set to three genes. The gene IDs from these clusters were extracted individually and tested for GO term representation using the functional annotations with TopGO (Rahnenfuhrer, 2022). To characterize further the expression differences of the ant DEGs, we investigated how the expression of the ant DEGs differed between the three parasite load categories (uninfected, lowly infected and highly infected). To do so, we ran three LRT tests (same syntax of full and reduced models as described before) on a gene subset that only contained the ant DEGs: one for each of the three possible combinations of two of the three parasite load categories. Furthermore, we performed KEGG enrichment analysis by comparing the KEGG IDs of the transcriptome to the ant DEGs within different clusters using the clusterProfiler package on R (Wu et al., 2021). Finally, the transcriptome was BLASTed to a protein database for all invertebrates (downloaded on 10/08/2021) using the blastx algorithm of diamond BLAST (Buchfink et al., 2014) with an e-value cut-off of 1e-5. All BLAST hits per gene with the lowest E-value were extracted to identify which proteins the transcripts are potentially encoding. The individual gene functions of these BLAST hits were studied in the literature and using UniProt.

We processed the raw RNAseq data from our cestode samples in the same way as for the ants, except that we filtered out contaminating sequences from *T. nylanderi, H. sapiens, E. coli* and adapters, and vectors using FastQScreen. Furthermore, we used a transcriptome assembly of *A. brevis* (Stoldt et al., 2021) to map reads and based on this, estimated gene abundances. We also analysed the cestode gene count matrix using DESeq2, with a likelihood ratio test (LRT) to compare a model that included parasite load (lowly infected and highly infected) and colony identity as factors, to a reduced model including only colony identity. This produced a list of cestode genes with expression differences associated with the parasite load (cestode DEGs). We divided the cestode DEGs into upregulated and downregulated in the high parasite load category. The cestode DEGs in these groups were diamond BLASTed using the same invertebrate database as used for the ant DEGs and functionally annotated using TopGO. Gene functions were once again studied in the literature using UniProt. Furthermore, similar to the ant DEGs, we performed enrichment analysis on the extracted KEGG IDs from EGGNOG mapper using clusterProfiler. Finally, to investigate whether some of those genes encode for proteins that are secreted into haemolymph of the host ants (Hartke et al., 2022), we BLASTed (BLAST v.2.9.0; Altschul et al. 1990) the cestode DEGs against the RNA sequences of secreated cestode proteins.

We also compared gene expression between uninfected ants from infected colonies and uninfected colonies. When we plotted a PCA for the uninfected samples we found one sample in the uninfected colonies to be an outlier (figure S2). However, considering that we found no further indication that this sample was problematic (e.g., in our laboratory notes, sequence quality control), we included it in the analysis. Here we used the colony infection status as the variable factor using a Wald test on DESeq2 for a pairwise comparison of gene expression between different colony infection status (whether the sample came from an infected colony or not). Similarly, to the cestode DEGs, we split up the colony infection DEGs into overexpressed and underexpressed genes in the infected category. The colony infection DEGs were functionally annotated using TopGO and clusterProfiler and were BLASTed using diamond BLAST.

### WGCNA analysis

We performed a weighted gene co-expression network analysis (WGCNA; Langfelder & Horvath, 2008) to investigate the effect of parasite load on gene networks. For this, we transformed the gene count data using variance stabilizing transformation from DESeq2. Networks were constructed using all the ant samples including the samples from uninfected colonies. WGCNA was performed with a soft threshold power of 5 and a minimum module size of 250. To test whether parasite load affected gene expression of the constructed networks, we correlated the module expression to parasite load using Pearson’s correlation and performed GO enrichment analysis on the significantly correlated modules.

## Results

### Influence of parasite load on morphology

Infected ant workers exhibited a smaller head width than uninfected ones (F= 33.442, p = 7.344e-09; Fig. 1a). Yet, worker head width did not further shift with increasing parasite load (F=0.536, p = 0.464; Fig. 1b). In contrast, the diameter of cestodes decreased with the number of cestodes in a single ant host (F=391.31, p = 2.2e-16; Fig. 1 c).

**Figure 1.**
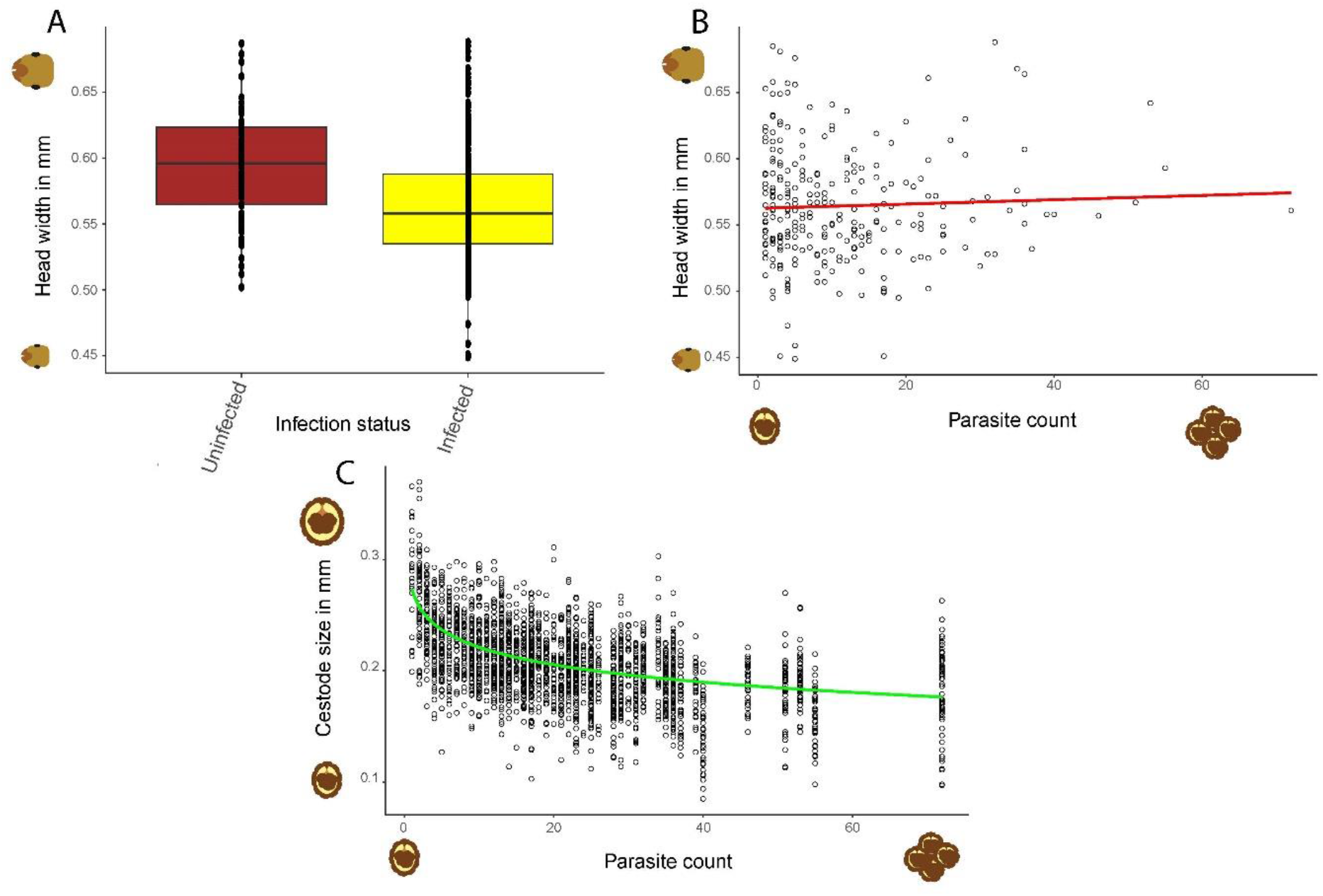
Changes in body size of parasites and hosts with parasite load. A) Infected workers had a smaller head width than uninfected workers (F= 33.442, p = 7.344e-09). B) No evidence for an association between head width of the host and parasite load (F=0.536, p = 0.464). C) Cestode size was negatively associated with parasite load (F=391.31, p = 2.2e-16).

### Influence of colony infection status on gene expression of uninfected workers

Uninfected workers from infected colonies differed in the expression of 46 genes in the fat body from those from uninfected colonies. Of these genes, 28 were upregulated in workers from uninfected colonies and 18 were upregulated in workers from infected colonies. Among the upregulated genes of workers from uninfected colonies, we found two significantly enriched GO terms represented by single genes, namely *transcription initiation by RNA polymerase* and *protein glycosylation*. The BLAST results revealed three homologues of probable cytochrome P450 6a14, which may have a function in insect hormone metabolism and synthetic pesticide degradation (Scanlan et al. 2020). For the overexpressed genes of workers from infected colonies, we found two enriched GO terms, also based on single genes *regulation of synaptic transmission, cholinergic* and *positive regulation of voltage-gated potassium channel activity*. However, the BLAST results provided limited information. Only three genes had annotations in *Drosophila melanogaste*r, two of which were related to brain function and the last one was a transmembrane and TPR repeat-containing protein CG4050 that transfers mannosyl residues to the hydroxyl group of serine or threonine residues (Larsen et al., 2017).

### Influence of parasite load on ant gene expression

We identified 122 differentially expressed genes (ant DEGs) in the fat body, whose expression varied according to infection status and/or parasite load. The cluster analysis grouped 120 of these ant DEGs into three clusters with different expression patterns. Cluster 1 consisted of 62 genes with high expression in highly infected workers, while the uninfected and lowly infected workers showed low expression of these genes (Fig. 2a). These genes were enriched for *oxidative stress response* (Fig. 2d). Pairwise comparisons revealed that 17 of these genes were differentially expressed between lowly and highly infected ants. In addition, ten more genes differed in expression between highly and uninfected workers than between lowly and uninfected workers (Table S3). Genes of note in this cluster include two homologues of genes encoding the dual-oxidase II (figure 3a and b). Both differed significantly between highly infected and uninfected workers, with low-infected workers not differing significantly from either. This protein mainly has been shown to play a role in the innate immunity, limiting microbial proliferation in *Drosophila melanogaster* guts (Ha et al., 2005; Lee et al., 2015). Other genes of note encode tyrosine-protein kinase Src64B, which may be involved in the development of neural tissue and smooth muscle (Simon et al., 1985) as well as salivary gland development (Sopko & Perrimon, 2013). Furthermore, there were three proteins involved in eye development, including protein tincar (Fig 3c; Boulton et al. 2000; Hirota et al. 2005; Rahman et al. 2012), which differed in expression between highly infected and uninfected workers, while there were no expression differences to the low-infected group in both cases.

**Figure 2.**
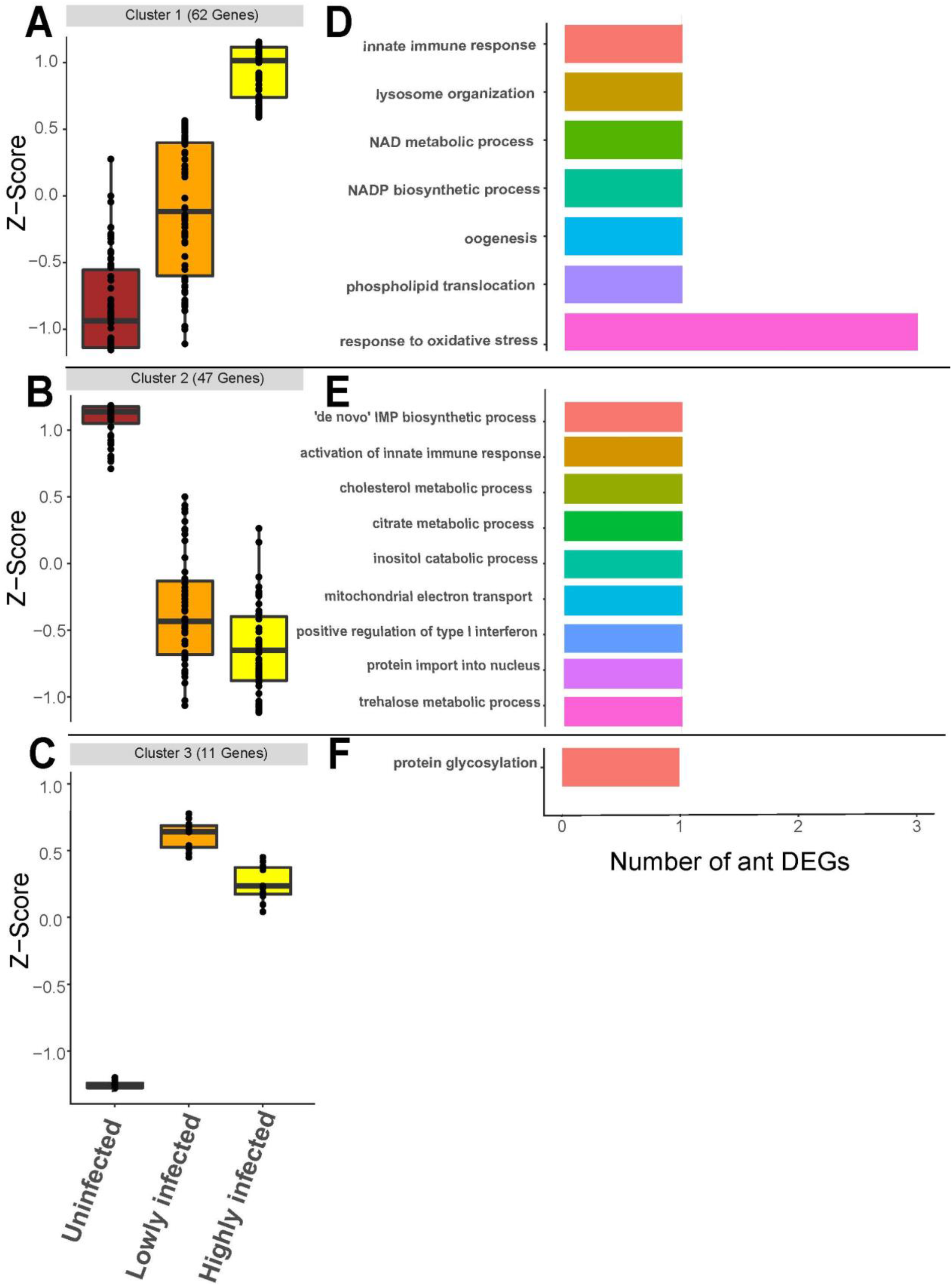
Clustering and visualization of 120 (out of 122) of the ant DEGs. The genes could be grouped into three clusters according to their expression patterns: A) Cluster 1, B) Cluster 2 and C) Cluster 3. GO enrichment analyses of these clusters (D for cluster 1, E for cluster 2 and F for cluster 3) were calculated with topGO using the weight01 algorithm and the GO annotations of the biological processes of the clusters were compared with the whole transcriptome using Fisher’s exact test. Each bar represents a significantly enriched GO term in each cluster, the x-axis represents the number of significant genes.

**Figure 3.**
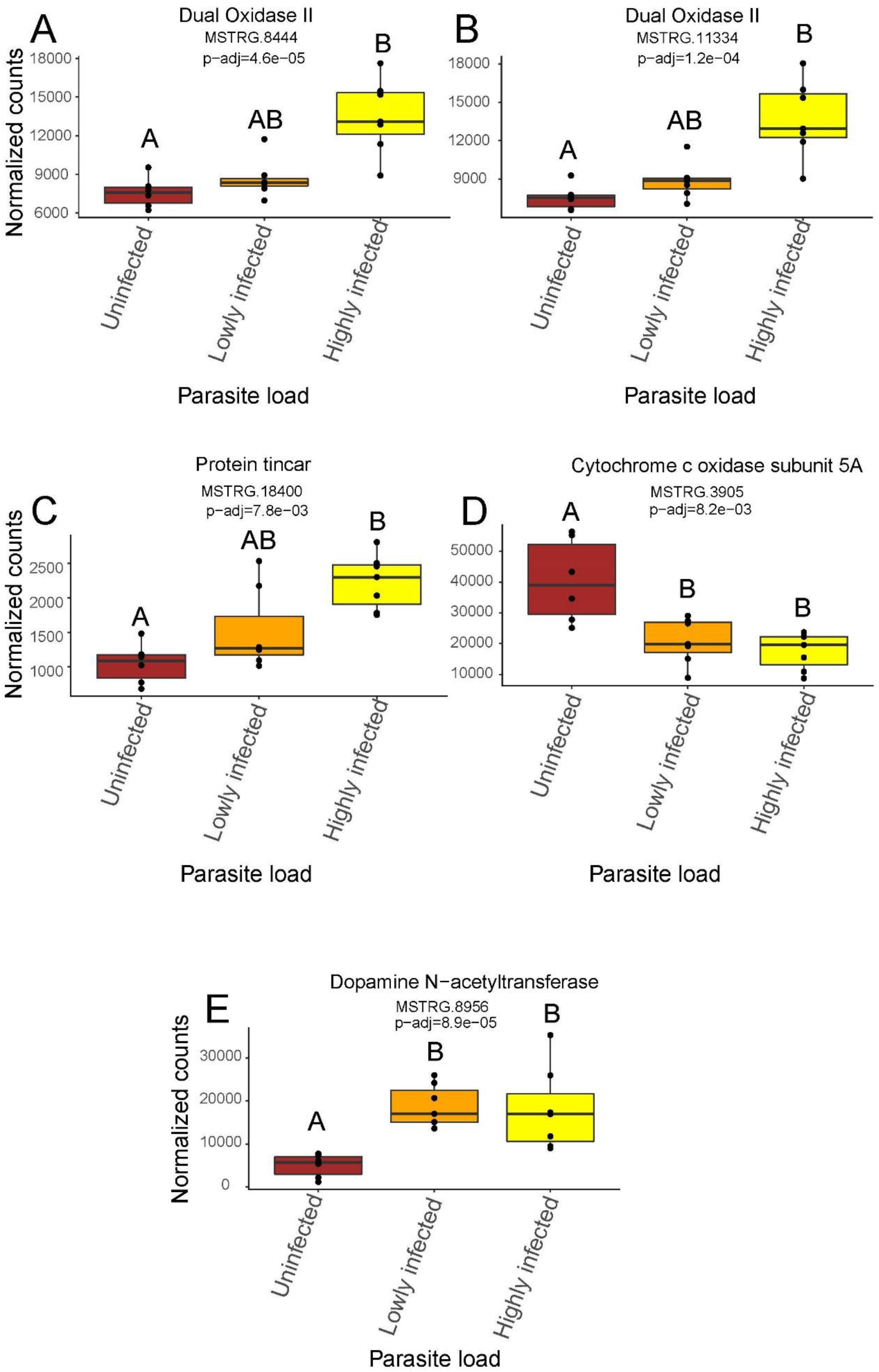
Selected ant DEGs with notable functions. A-C shows DEGs from cluster 1, which were upregulated in highly infected individuals (but not in lowly infected ones) compared to the uninfected individuals. D belonging to cluster 2, upregulated in uninfected workers. E belonging to cluster 3, with DEGs downregulated in uninfected workers.

The second cluster included 47 DEGs (Fig. 2b) and as the graphical representation indicates that these genes were generally upregulated in uninfected workers and downregulated in both groups of infected workers. Nevertheless, pairwise comparisons revealed fourteen genes that differed in expression between lowly and highly infected workers and nine additional DEGs between highly and uninfected workers that did not differ between uninfected and lowly infected workers (Table S3). Among the enriched GO terms of this cluster, we found *activation of innate immune response*. From the BLAST results, genes of interest include those encoding the cytochrome c oxidase subunit 5A (Fig. 3d), which is the last enzyme in the mitochondrial electron transport chain, driving oxidative phosphorylation (Hartley et al., 2019) and therefore plays an important role in energy metabolism.

Finally, cluster 3 contained 11 DEGs, which were downregulated in the uninfected workers but upregulated in the lowly and highly infected workers (Fig. 2c). For none of these genes differed in expression between the lowly and highly infected group (Table S3). Only one GO term was found to be enriched, which was *protein glycosylation*, based on a single gene. Genes of interest include one encoding a dopamine N-acetyltransferase (Fig. 3e), which regulates sleep homeostasis by limiting the accumulation of serotonin (Brodbeck et al., 1998; Davla et al., 2020; Shaw et al., 2000).

We found no significantly enriched KEGG IDs for any of the clusters. In the WGCNA analyses, we identified a total of 10 modules with a minimum module size of 100. However, none of these clusters were significantly associated with parasite load (figure S3).

### Influence of parasite load on cestode gene expression

Analysis of the cestode dataset revealed 377 DEGs between cestodes from highly and lowly infected ants (cestode DEGs), of which 357 were upregulated in cestodes co-infecting a single host with many others.

Ten highly expressed genes in this group are involved in the serine/glycine interconversion pathway. Serine proteinases are associated with multiple functions in cestodes, including defence against the host immune system (Izvekova et al., 2021; Patel, 2017) and lipid metabolism (Webb & Mettrick, 1975). In addition, we found overexpression of mitochondrial uncoupling protein 4, which is involved in the response to starvation in *C. elegans* (Figure 4a) and regulates the long-chain fatty acid metabolism, primarily in response to nutrient availability. The production of these lipid-signalling proteins regulates several processes that may control lifespan and adaptation to starvation (O’Rourke et al., 2013). Finally, we found an upregulation of a gene encoding a vesicular glutamate transporter, which in *C. elegans* is required for the detection of preferred food sources (Harris et al., 2014). The GO enrichment analysis (Fig. 4c) of these upregulated genes in parasites from highly infected hosts revealed transport functions, among which are *carbohydrate transmembrane transport, regulation of pH* and *sodium ion transport*. When comparing these genes to those proteins released into the haemolymph of host ants (Hartke et al. 2022) using BLAST, we found three matching genes, none of which were annotated.

**Figure 4.**
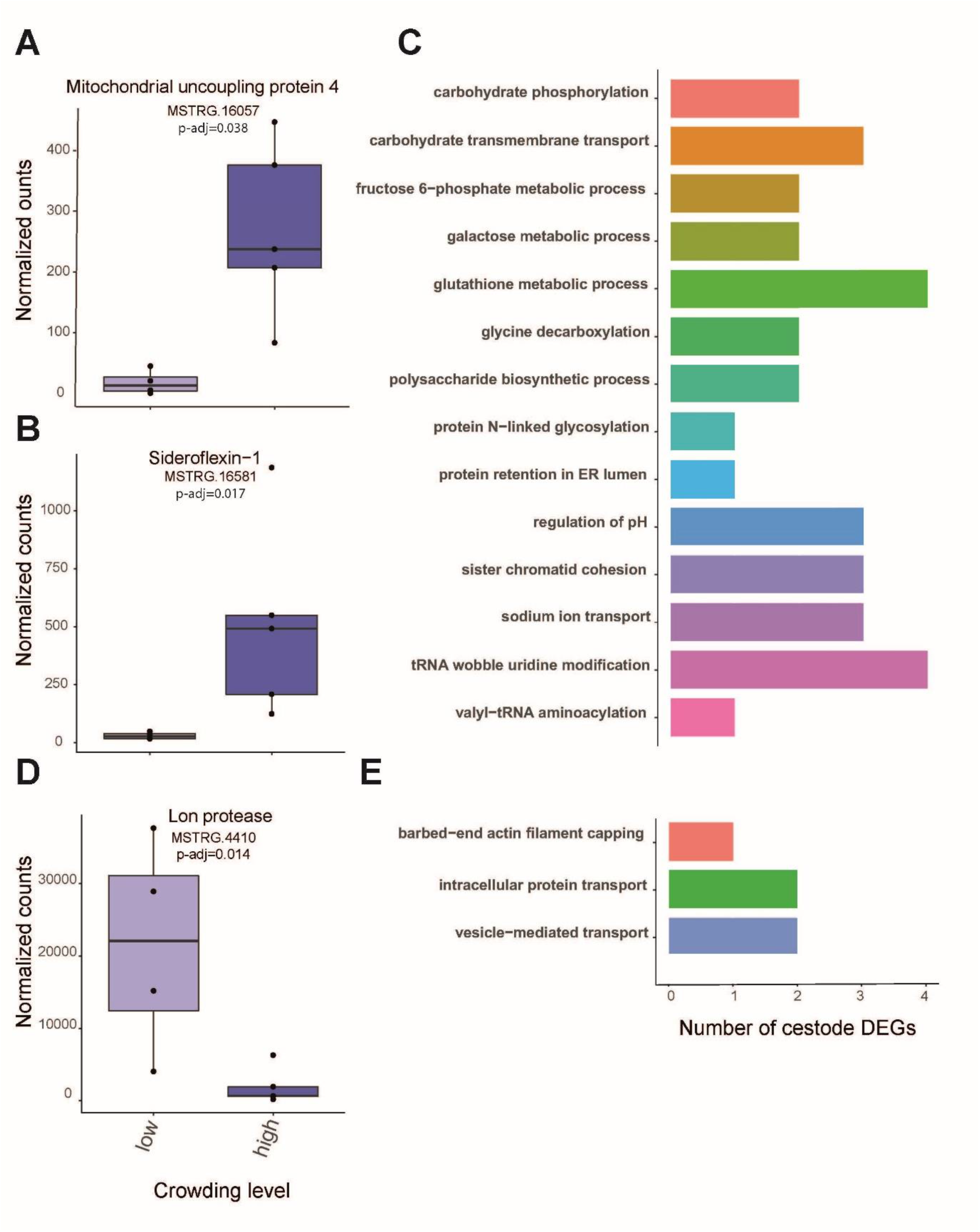
Changes in gene expression in cestodes from highly infected workers. A-C: Expression differences for three selected cestode DEGs. D. Barplots representing the GO terms for the genes that were upregulated with higher parasite load. E. Barplots representing the GO terms for the genes that were upregulated with lower parasite load.

Among the 20 genes upregulated in parasites residing alone or with one other parasite in a host, we found a gene encoding a Lon protease (Fig. 4d), which is a serine-dependent protease that ensures the maintenance of the integrity of the mitochondrial genome (Kaunisto & Suhonen, 2013; Nargund et al., 2012). In the GO enrichment of upregulated genes in cestodes from lowly infected hosts, we found *vesicle-mediated transport* as a significantly enriched function, as well as *carbohydrate transmembrane transport* (both 2 genes) and fructose 6-phosphate metabolic process (Fig. 4e), both involved in the absorption of nutrients. No significantly enriched KEGG IDs were found in the entire dataset. Finally, we detected one BLAST match with a secreted cestode protein in the DEGs, but again the gene had no annotation.

## Discussion

The aim of our study was to investigate whether parasite load by the cestode *A. brevis* affects gene expression and morphology of *T. nylanderi* ant workers. About a quarter of all 120 clustered differentially expressed genes in the host fat body responded to parasite load, and in heavily infected workers, genes responsive to oxidative stress function were particularly upregulated. The cestodes were also strongly influenced by crowding. Our transcriptomic results indicate that co-infecting parasites suffer from poorer access to nutrients as reflected by their smaller body size.

### Influence of parasite load on host gene expression

Both pairwise comparisons as well as clustering of ant DEGs revealed that the expression of many genes responded to parasite load, whereas others responded to infection in an all-or-nothing manner such as those DEGs grouped in cluster 3. Looking at the GO enrichment of cluster 1, the most enriched term was *oxidative stress response*. A higher parasite load could be responsible for more reactive oxygen species, i.e., more oxidative stress, as has been shown in rodents infected with a high load of the protozoan parasite *Trypanosoma cruzi* (Vilar-Pereira et al., 2021) or in erythrocytes infected with malaria (Becker et al., 2004). However, increased response to oxidative stress could also be a way to extend lifespan (Ristow & Schmeisser, 2011). Infected *T. nylanderi* host workers live far longer than their uninfected nestmates (Beros et al., 2021), and this may be explained in part by a stronger response to oxidative stress. In our system, the parasite might benefit from the host’s life extension, as this increases the chances that the final host, a woodpecker, will prey upon the infected ant. None of the genes in clusters 2 and 3 were involved in the regulation of oxidative stress. Based on the expression plots (Fig. 2) and the pairwise comparisons, we interpret that the enrichment detected in cluster 1 is mainly due to the large number of DEGs that differ between uninfected and lowly infected workers on the one hand and highly infected workers on the other, whereas hardly any differences were found between uninfected and lowly infected workers. This would mean either that only a high parasite load may trigger oxidative stress in the host, to which the ant responds by upregulating the response genes, or that only many parasites jointly can influence the host to upregulate oxidative stress genes and thus live longer. To date, it has not been studied how parasite load relates to the longevity of individual workers, and we plan to investigate this. We would argue that it is more likely that the oxidative stress response is a side-effect of parasite load or a consequence of heightened immune response of the host, as higher parasite burden has been linked to a high oxidative stress response in other parasite-host systems (De Coster et al., 2012; Stocker et al., 1985). The higher stress from infection is also supported by the differential innate immune response, which we find confirmed in the GO term analysis.

While examining the functions of differentially expressed genes, a gene of interest was one encoding *dual oxidase II*, two homologues of which were upregulated in highly infected workers (but not in lowly infected ones) compared to uninfected workers. *Dual oxidase II* is involved in limiting microbial proliferation in the gut of *D. melanogaster* flies (Ha et al., 2005; Lee et al., 2015), suggesting possible changes in the gut microbiome of highly infected workers. In general, the gut microbiome of *T. nylanderi* appears to be colony-dependent, with little difference between castes, although colonies with a greater microbial diversity are more productive (Segers et al., 2019). A difference in the gut microbiome caused by cestode infection might be linked to parasite load, and this should be investigated further.

Confirming previous results (Feldmeyer et al., 2016; Stoldt et al., 2021), we identified an enrichment for innate immune response in cluster 2, containing genes with high expression in uninfected workers. However, we should note that this GO enrichment was based on a single gene. Another notable difference between infected and uninfected ants is the number of differentially expressed genes involved in transport processes. Examples of enriched GO terms include *mitochondrial electron transport* and *protein import into the nucleus*, both represented by a single gene. In addition, we found genes encoding proteins with transport functions, such as cytochrome c oxidase subunit 5A (Hartley et al., 2019) and cystinosin homologues isoform X3 (Yu et al., 2008). Differential expression of genes involved in transport is frequently observed during parasite infection (Berger et al., 2021; Kremer et al., 2013). Transport genes are thought to be involved in cestode-host communication, and more commonly as a means of manipulating host phenotypes (Berger et al., 2021). Thus, the upregulation of transport genes may indicate active manipulation by *A. brevis*, which should be investigated further, even though we found no evidence in cluster 1 that heavily infected ants particularly upregulate transport genes. A final candidate gene in cluster 3 encodes a dopamine N-acetyltransferase gene involved in sleep regulation in insects (Brodbeck et al., 1998; Davla et al., 2020; Shaw et al., 2000). Its upregulation in lightly and heavily infected workers could possibly explain why infected ants are so inactive. Biogenic amines such as dopamine are important in the brain for regulating behaviour. So, if the parasite is interfering with host gene expression in the fat body, it is probably doing so in the wrong tissue. However, it has already been shown in other parasite systems that parasites do not always exclusively influence the physiology of tissues they are located in (Laothamatas et al., 2003). Inhibition of this gene using RNAi may help to understand whether it indeed plays a role in the behavioural changes of infected animals (Berger et al., 2021; Kremer et al., 2013).

Parasite load was unrelated to host body size, which is in contrast to previous results demonstrating a negative correlation between ant head size and parasite load (Scharf et al., 2012). The discrepancy may be due to different methodologies: Scharf et al. (2012) measured the entire head surface, while we used only head width a common measure of body size in ants. It is possible that increasing parasite load causes a shortening of the head but does not cause a reduction in head width from eye to eye. This might be related to a reported atrophy of the mandible closer muscle in infected ants (Feldmeyer et al., 2016). Changes in head shape have not been studied in relation to parasite load, but is a known consequence of parasite infection in various systems (Johannessen, 1973; Johnson & Sutherland, 2003). Our measurements did demonstrate a clear reduction in head width related to the cestode infection *per se*. This would mean that head width falls into our second category of phenotypic changes, those that shift with infection in an on/off manner.

### Influence of crowding on cestode morphology and gene expression

We can show that crowding (multiple cestodes in a single host) leads to reduced body size in cysticercoid larvae of our parasitic cestode. This has been previously noted in *A. brevis* (Scharf et al., 2012) and several other cestode species (Benesh, 2019; Heins et al., 2002) and may indicate reduced nutrient availability for cestodes residing in heavily infected workers. This was also evident in the transcriptional shifts observed with crowding. When looking at the functional annotation of the cestode DEGs, many functions related to transport become apparent, similar to the ant dataset. For parasites living under crowded conditions, these include *carbohydrate transmembrane transport* (3 genes), *pH regulation* (3 genes) and *sodium ion transport* (3 genes). These enriched GO terms could indicate the need to transport in nutrients and salts and to regulate the pH. In contrast, parasites infecting an ant host alone upregulated DEGs enriched for *intracellular protein transport* (2 genes) and *vesicle-mediated transport* (2 genes). Cestodes are known to release vesicles into their environment that contain miRNAs (Ancarola et al., 2020, 2017). In our study system, tetraspanin, a protein considered indicative of extracellular vesicles, was identified among the cestode proteins released into ant haemolymph, suggesting that *A. brevis* uses this pathway to interact with its host (Hartke et al. 2022). A function of extracellular vesicles in parasite-host communication or manipulation has been suggested also in other systems (Coakley, et al., 2015; 2016; Siles-Lucas et al., 2015; Tritten & Geary, 2018), this all points to a need of our parasites living alone to communicate more with (or manipulate) their hosts. A GO term supported by four genes for cestodes living under crowded condition was glutathione metabolic process. An upregulation in glutathione provides protection from and tolerance of severe stress (Maher, 2005), again indicating physiological stress. Ten serine proteases that are upregulated in cestodes living under crowded conditions provide more insights. Among other things, serine proteases are related to the ability of cestodes to evade the host immune system (Izvekova et al., 2021; Patel, 2017). Since infection with multiple parasites could trigger a stronger host immune response, this enhanced evasion of the host immune system might be necessary. Under crowded conditions, there may also be competition for food among parasites. Indeed, we find evidence for this. For example, mitochondrial uncoupling protein is upregulated in co-infecting cestodes, which has been shown to allow at least *C. elegans* to adapt to food deprivation (Folick et al., 2015; Lapierre et al., 2011; O’Rourke et al., 2013; Ramachandran et al., 2019). Although these effects could generally be considered negative for the parasite, nutrient stress fits surprisingly well with the oxidative stress experienced by the host during high parasite loads, as oxidative stress is inherently life-prolonging (Ristow & Schmeisser, 2011). This is mainly because low food intake has been shown to prolong life in both fungi (Lin et al., 2000) and animals (Mathis et al., 2017). In this context, it should not be forgotten that the parasite must live for a very long time as well; it is not enough to increase the life expectancy of the host to several years; the cysticercoid larvae also have to stay alive until they reach their final host. If they do so and have lived with other cestodes in the intermediate host, they are also likely to co-infect their final avian host, which could lead to a reduction in fitness during their reproductive phase, although a fitness reduction has not been shown yet because of crowding (Benesh, 2019). The consequences of crowding are thus often manifold and lasting.

## Conclusion

Not only the infection *per se* but also the parasite load can cause multiple consequences in the physiology for both the host and the parasite. In the host of our study system, we found that many of the differentially expressed genes respond to parasite load. Gene functions were indicative of oxidative stress and innate immunity, suggesting general stress and possibly also extended lifespans in highly infected hosts. We also find evidence that sole parasites invest more in vesicular transport mechanisms. On the other hand, crowded parasites seem to suffer from nutrient stress and invest more in the evasion of the hosts’ immune system, nutrient transport, and stress resistance. These deeper insights into the host-parasite interaction have demonstrated that both the parasite and the host are affected by changes in parasite load.

## Acknowledgements

We would like to thank Jenny Fuchs for creating the illustrations of *T. nylanderi* and *A. brevis*, as well as designing Figure S1. Furthermore, thanks go to Marina Choppin for her help with the dissection setup and Marion Kever for doing some of the RNA isolations needed for this study as well as her instructions on how to do them. Finally, we would like to thank Falk Butter and Peter Baumann for their input and fruitful discussions regarding the data analysis. This project was funded by the Deutsche Forschungsgemeinschaft (DFG, German ResearchFoundation) – GRK2526/1 – Projectnr. 407023052.

## Supplementary materials

**Figure S1.**
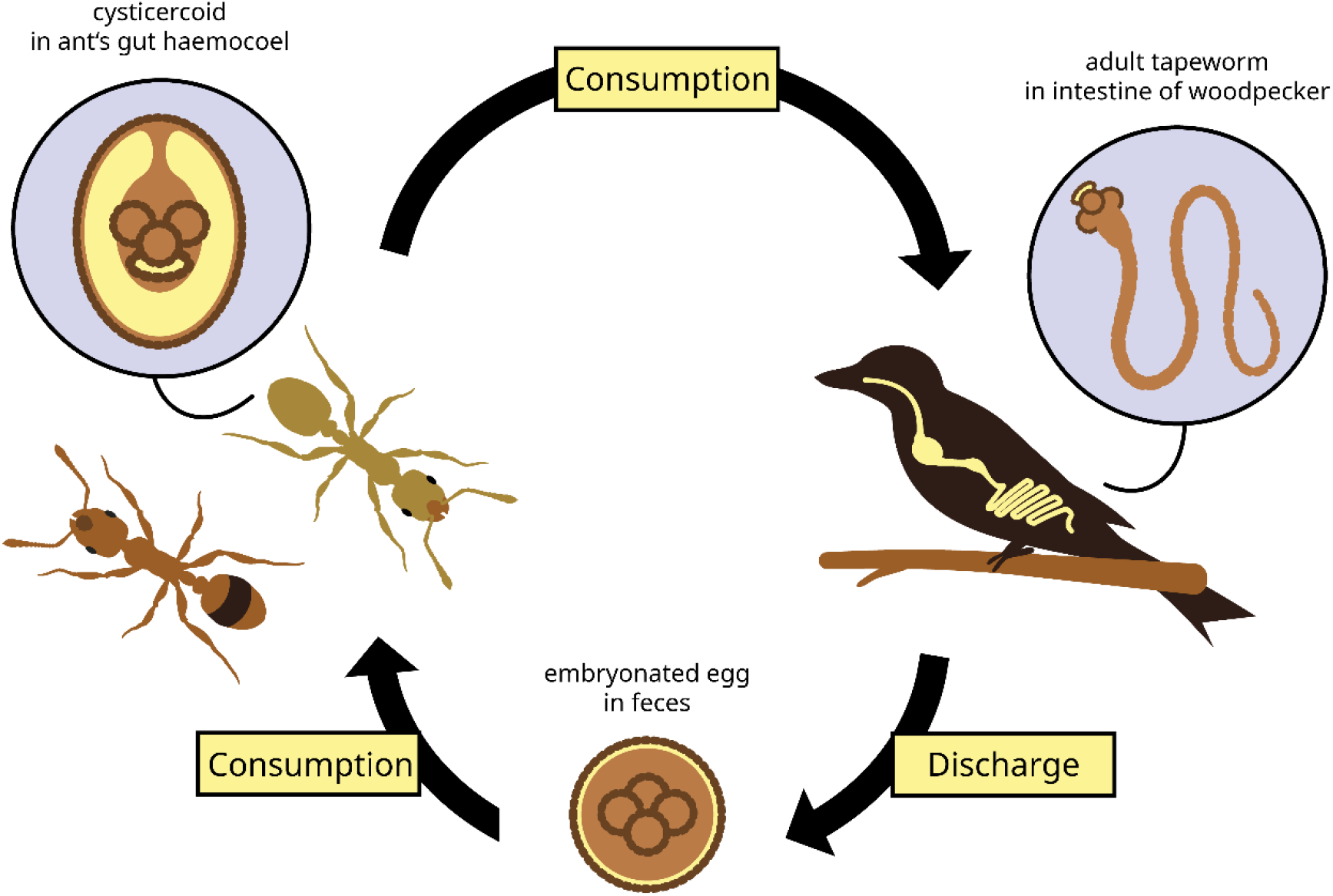
A visualisation of the life cycle of *Anomotaenia brevis*. Their eggs are located in the discharge of woodpeckers which is consumed by *Temnothorax nylanderi*, where *A. brevis* spends its cysticercoid stage. When the ants are then consumed by woodpeckers they grow into their adult life stage.

**Figure S2.**
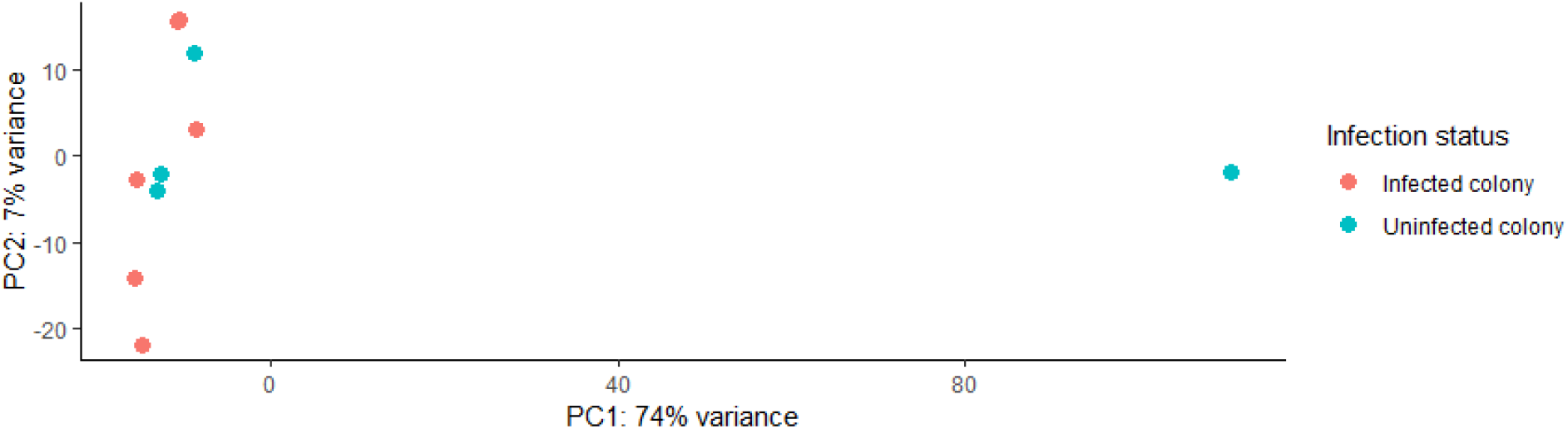
PCA of the uninfected colonies where the colour represents infected (red) or uninfected (blue) colonies. No clustering based on infection status was observed in the data, however, one uninfected sample played a large role in PC1, which explains 74% of the variance. This sample was however not removed due to a lack of evidence that anything had happened to this sample or that anything had gone wrong during sequencing.

**Figure S3.**
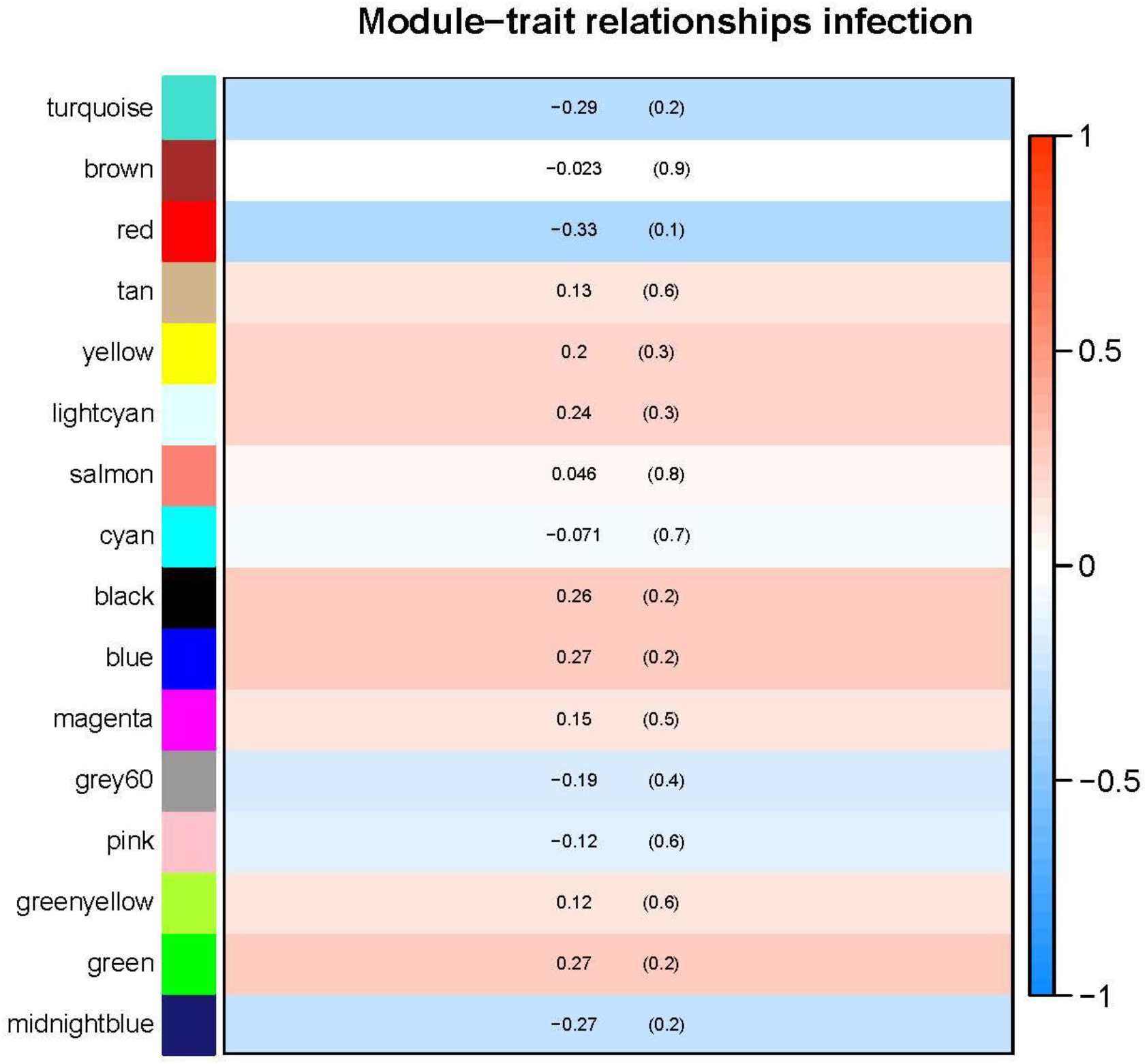
Heatmap of the different WGCNA modules with their correlation coefficient and significance. None of the modules correlate significantly with parasite load.

**Table S1:** Overview of the colonies used for the analysis. The colonies marked in bold were used for gene expression analysis of the ants. From colonies indicated by an asterisk, we extracted the RNA from cestodes as well

| Colony identity | Year of collection | Number of queens | Total number of workers | Number of infected workers | Infection rate | Uninfected workers dissected |
| --- | --- | --- | --- | --- | --- | --- |
| LBW20TS15 | 2020 | 1 | 77 | 31 | 0.40 | 7 |
| **LBW20TS49\*** | 2020 | 1 | 65 | 4 | 0.06 | 3 |
| LBW20TS52* | 2020 | 1 | 44 | 16 | 0.36 | 4 |
| **LBW20TS58\*** | 2019 | 1 | 164 | 16 | 0.10 | 5 |
| LBW20TS61 | 2019 | 1 | 89 | 1 | 0.01 | 2 |
| LBW20TS86 | 2019 | 1 | 65 | 2 | 0.03 | 1 |
| **LBW20TS89** | 2019 | 1 | 93 | 5 | 0.05 | 2 |
| LBW20TS90 | 2019 | 1 | 129 | 3 | 0.02 | 2 |
| LBW20TS97 | 2020 | 1 | 196 | 3 | 0.01 | 2 |
| **LBW20TS106\*** | 2020 | 1 | 72 | 16 | 0.22 | 4 |
| LBW20TS107 | 2020 | 1 | 137 | 4 | 0.03 | 2 |
| LBW20TS108 | 2020 | 1 | 177 | 126 | 0.71 | 26 |
| **LBW20TS113\*** | 2020 | 1 | 121 | 24 | 0.20 | 4 |
| Lbw20TS117 | 2020 | 1 | 62 | 1 | 0.02 | 2 |
| LBW20TS118 | 2020 | 1 | 86 | 1 | 0.01 | 2 |
| LBW20TS120 | 2020 | 1 | 16 | 2 | 0.13 | 2 |
| LBW20TS121 | 2020 | 1 | 72 | 8 | 0.11 | 2 |
| LBW20TS132 | 2020 | 1 | 88 | 1 | 0.01 | 2 |
| **LBW20TS145** | 2020 | 1 | 17 | 2 | 0.12 | 2 |
| **LBW20TS149** | 2020 | 1 | 76 | 8 | 0.11 | 2 |
| LBW20TS153 | 2020 | 1 | 66 | 1 | 0.02 | 2 |
| **LBW20TS9** | 2020 | 1 | 68 | 0 | 0 | 5 |
| **LBW20TS21** | 2020 | 1 | 167 | 0 | 0 | 5 |
| **LBW20TS99** | 2019 | 1 | 99 | 0 | 0 | 5 |
| **LBW20TS105** | 2020 | 1 | 49 | 0 | 0 | 5 |

**Table S2.** Overview of all samples used for RNA seq analysis and time and day of sampling. Samples marked with an asterisk are cestodes from ants represented in the RNA seq analysis for the ants and vice versa. The sample marked in grey (C16) was removed from analysis due to poor read quality.

**Highly infected workers**
| Code | Colony | Infected | Date | Time | Parasite count |
| --- | --- | --- | --- | --- | --- |
| J2* | LBW20TS106 | y | 30-7-2020 | 18:40 | 23 |
| M19* | LBW20TS113 | y | 4-8-2020 | 10:30 | 18 |
| S1 | LBW20TS145 | y | 31-7-2020 | 15:10 | 34 |
| T3 | LBW20TS149 | y | 6-8-2020 | 13:15 | 28 |
| D16* | LBW20TS58 | y | 31-7-2020 | 14:00 | 51 |
| G5 | LBW20TS89 | y | 4-8-2020 | 10:05 | 31 |
| B5* | LBW20TS49 | y | 28-7-2020 | 15:00 | 29 |

**Lowly infected workers**
| Code | Colony | Infected | Date | Time | Parasite count |
| --- | --- | --- | --- | --- | --- |
| J10 | LBW20TS106 | y | 4-8-2020 | 13:45 | 1 |
| M7 | LBW20TS113 | y | 29-7-2020 | 13:25 | 1 |
| S2 | LBW20TS145 | y | 7-8-2020 | 09:20 | 3 |
| T10 | LBW20TS149 | y | 29-7-2020 | 18:30 | 5 |
| D15 | LBW20TS58 | y | 4-8-2020 | 09:50 | 2 |
| G7 | LBW20TS89 | y | 5-8-2020 | 13:15 | 6 |
| B3 | LBW20TS49 | y | 24-7-2020 | 14:00 | 2 |

**Uninfected workers from infected colonies**
| Code | Colony | Infected | Date | Time | Parasite count |
| --- | --- | --- | --- | --- | --- |
| J19 | LBW20TS106 | n | 3-8-2020 | 19:00 | 0 |
| M24 | LBW20TS113 | n | 31-7-2020 | 16:35 | 0 |
| S5 | LBW20TS145 | n | 6-8-2020 | 10:40 | 0 |
| T11 | LBW20TS149 | n | 6-8-2020 | 15:50 | 0 |
| D20 | LBW20TS58 | n | 7-8-2020 | 11:40 | 0 |
| GX | LBW20TS89 | n | 10-8-2020 | 17:50 | 0 |
| B6 | LBW20TS49 | n | 7-8-2020 | 15:00 | 0 |

**Uninfected colonies**
| Code | Colony | Infected | Date | Time | Parasite count |
| --- | --- | --- | --- | --- | --- |
| V1 | LBW20TS9 | n | 5-8-2020 | 13:05 | 0 |
| W3 | LBW20TS21 | n | 4-8-2020 | 15:25 | 0 |
| X4 | LBW20TS99 | n | 10-8-2020 | 11:35 | 0 |
| Y2 | LBW20TS105 | n | 6-8-2020 | 17:35 | 0 |

**Cestodes**
| Code | Colony | Infected | Date | Time | Parasite count |
| --- | --- | --- | --- | --- | --- |
| B4 | LBW20TS49 | y | 21-7-2020 | 17:55 | 1 |
| B5* | LBW20TS49 | y | 28-7-2020 | 15:00 | 29 |
| C16 | LBW20TS52 | y | 4-8-2020 | 19:15 | 1 |
| C9 | LBW20TS52 | y | 31-7-2020 | 09:30 | 22 |
| D15 | LBW20TS58 | y | 10-08-2020 | 09:25 | 2 |
| D16* | LBW20TS58 | y | 31-7-2020 | 14:00 | 51 |
| J10 | LBW20TS106 | y | 4-8-2020 | 13:45 | 1 |
| J2* | LBW20TS106 | y | 30-7-2020 | 18:40 | 23 |
| M11 | LBW20TS113 | y | 5-8-2020 | 09:20 | 1 |
| M19* | LBW20TS113 | y | 4-8-2020 | 10:30 | 18 |

**Table S3.** Overview of the pairwise comparisons between ant DEGs

| Cluster ID | No. of genes in cluster | Uninfected v lowly infected | Uninfected vs highly infected | Lowly infected vs highly infected |
| --- | --- | --- | --- | --- |
| Cluster 1 | 62 | 33 (53%) | 43 (69%) | 17 (27%) |
| Cluster 2 | 47 | 38 (81%) | 47 (100%) | 14 (30%) |
| Cluster 3 | 11 | 11 (100%) | 9 (82%) | 0 (0%) |

